# Evidence and urgency related EEG signals during dynamic decision-making in humans

**DOI:** 10.1101/2020.10.02.323683

**Authors:** Y. Yau, T. Hinault, M. Taylor, P. Cisek, L.K. Fellows, A. Dagher

## Abstract

A successful class of models link decision-making to brain signals by assuming that evidence accumulates to a decision threshold. These evidence accumulation models have identified neuronal activity that appears to reflect sensory evidence and decision variables that drive behavior. More recently, an additional evidence-independent and time-variant signal, named urgency, has been hypothesized to accelerate decisions in the face of insufficient evidence. However, most decision-making paradigms tested with fMRI or EEG in humans have not been designed to disentangle evidence accumulation from urgency. Here we use a face-morphing decision-making task in combination with EEG and a hierarchical Bayesian model to identify neural signals related to sensory and decision variables, and to test the urgency-gating model. We find that an evoked potential time-locked to the decision, the centroparietal positivity, reflects the decision variable from the computational model. We further show that the unfolding of this signal throughout the decision process best reflects the product of sensory evidence and an evidence-independent urgency signal. Urgency varied across subjects, suggesting that it may represent an individual trait. Our results show that it is possible to use EEG to distinguish neural signals related to sensory evidence accumulation, decision variables, and urgency. These mechanisms expose principles of cognitive function in general and may have applications to the study of pathological decision-making as in impulse control and addictive disorders.

**Significance Statement:** Perceptual decisions are often described by a class of models that assumes sensory evidence accumulates gradually over time until a decision threshold is reached. In the present study, we demonstrate that an additional urgency signal impacts how decisions are formed. This endogenous signal encourages one to respond as time elapses. We found that neural decision signals measured by EEG reflect the product of sensory evidence and an evidence-independent urgency signal. A nuanced understanding of human decisions, and the neural mechanisms that support it, can improve decision-making in many situations and potentially ameliorate dysfunction when it has gone awry.

## Introduction

Studies of decision-making typically assume a stochastic accumulation of sensory evidence with a decision being made once a threshold is reached (Gold and Shadlen, 2007; Ratcliff and McKoon, 2008). Drift diffusion models (DDMs) have a rich history and have successfully identified neuronal signals encoding evidence accumulation and decision threshold in simple, well-controlled experimental paradigms (Shadlen and Newsome, 2001; Kiani and Shadlen, 2009). However, interactive behavior is also determined by constantly changing and unpredictable environments. The notion that sensory evidence must achieve a critical threshold before the decision is made is difficult to reconcile with situations in which choices are made under temporal pressure or based on little to no sensory evidence. In such situations, something other than sensory evidence accumulation must contribute to choice commitment.

Convergent lines of research now support the notion of an additional “urgency signal” that non-selectively elevates activity towards unchanged action thresholds, such that less sensory evidence is required for decision commitment as time elapses (Cisek et al., 2009; Standage et al., 2011; Thura and Cisek, 2014; Murphy et al., 2016; Malhotra et al., 2018; Palestro et al., 2018). The level or urgency is thought to be stable in an individual although it will differ for different contexts (e.g. favoring speed versus accuracy) (Thura and Cisek, 2014; Berret et al., 2018; Reppert et al., 2018). It may be linked to phenotypical personality traits such as impulsivity (Carland et al., 2019). Primate single-cell recording studies indicate that this evidence-independent influence on the decision process is observable in the activity of neurons that reflect evolving decision formation (Ditterich, 2006; Churchland et al., 2008; Heitz and Schall, 2012; Hanks et al., 2014; Thura and Cisek, 2016).

Recently, this line of work has been extended to human decision-making using scalp electroencephalography (EEG). One outcome has been to identify separable EEG signals related to sensory evidence accumulation, decision variable, and motor response (O’Connell et al., 2012; Kelly and O’Connell, 2013; van Vugt et al., 2019). An event-related potential (ERP) labelled centroparietal positivity (CPP) appears to trace evidence accumulation (O’Connell et al., 2012). The urgency gating model (UGM) would predict that the CPP, as a reflection of the evolving decision variable, should reflect a combination of urgency and sensory evidence. However, in one study using a fixed visual stimulus, speed pressure was found not to affect the CPP (Steinemann et al., 2018). It may be that sudden-onset and discrete trial presentation coupled with short trial times often favoured in ERP may partly obscure the dynamics of an unfolding decision process. Attempts to use functional magnetic resonance imaging (fMRI) with evidence accumulation paradigms have provided evidence that a basal ganglia based urgency signal exists in humans (Nagano-Saito et al., 2012; Yau et al., 2020), as shown in primates (Thura and Cisek, 2017), but these lack the temporal resolution to precisely resolve urgency signalling.

In order to disambiguate sensory processing, decisional evidence accumulation, and urgency, we designed a dynamic decision-making task with a slowly morphing stimulus consisting of faces whose emotions transitioned from neutral to happy or sad (Yau et al., 2020). We exploited the high temporal resolution of EEG to tease apart neural determinants of human decision formation. First, we hypothesized that onset-locking the EEG to the start of the trial would allow us to detect signals related to facial emotion processing, sensory evidence accumulation, or setting of the decision threshold (i.e., N170 or P300). Second, by response-locking the EEG to the decision point, we sought to identify a neural signal (i.e., CPP) that ramps up in time and reflects the decision variable (or rate of evidence accumulation). Finally, by including easy and ambiguous trials, we aimed to identify an urgency signal associated with early responses despite ambiguous sensory evidence. Given the relatively large sample of subjects, and based on our previous work with this task, we investigated whether individuals may exhibit differing trait levels of the endogenous urgency signal.

## Methods

### Participants

74 right-handed young healthy adults (34 males; mean age 23.4 years ± 5.2 standard deviation) participated in the experiment for monetary compensation. All subjects gave informed consent prior to data acquisition and were screened for current or past diagnosis of a psychiatric disorder, neurological disorder, or concussion, and moderate to severe depression (score >5 on the Beck Depression Inventory (Beck et al., 1961)). The study was approved by the Montreal Neurological Institute Research Ethics Board.

### Experimental Design

Participants viewed short videos of a face “morphing” between expressions (Fig. 1). Each trial was preceded by a time-jittered fixation cross. Trials always began with a neutral expression and gradually transitioned into either a happy or sad emotion. Participants were instructed to predict whether the facial expression would be happy or sad by the end of the trial using their index and middle finger, respectively, via a button box in their right hand, and to respond whenever they felt confident enough to do so. Subjects were asked to respond both as quickly and as accurately as possible. Face stimuli were derived from the NimStim database (Tottenham et al., 2009) and manipulated using the STOIK MorphMan software (http://www.stoik.com/) to generate 18 intermediate faces that gradually transitioned in intensity of emotional expressions between a model’s neutral and happy or sad emotion. Thus, emotion levels varied from 0 to 19 in both directions. Trials lasted for a maximum of 60 frames over 6secs (i.e., 10 frames per sec), plus a final image of the correct emotion (with the emotion level >15) for 1sec either immediately after a response was made or at the end of the trial if the participant had not yet made a response. In the latter case, subjects could still respond during the final frame (i.e., 61^st^ frame) only if they had not yet done so earlier in the trial. Responses during this period were recorded but the frame would not change and persisted until the original 1sec window was over.

**Fig. 1.**
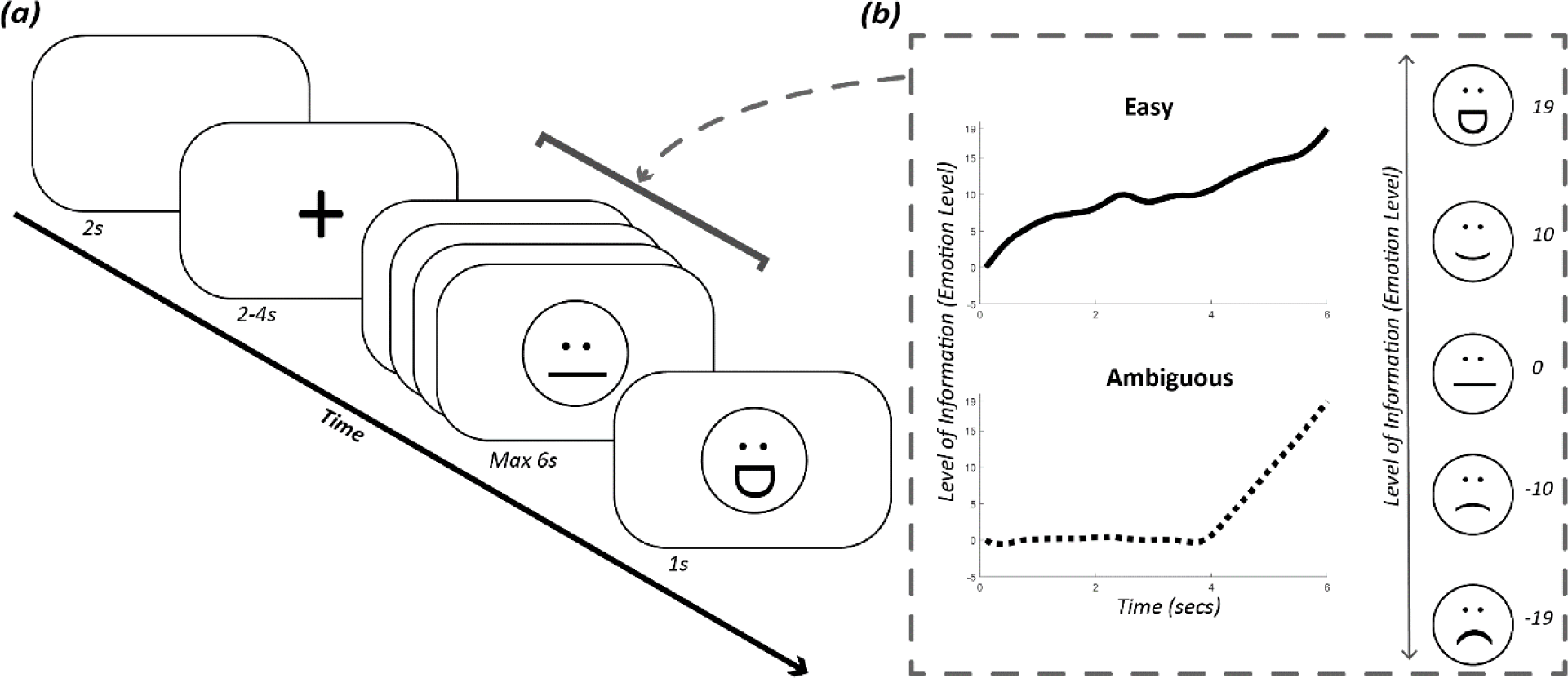
Schematic of task design. **(a)** Progression of a single trial begins with a blank image followed by a time-jittered fixation cross. A short video that start at a neutral facial expression which transitions into a happy or sad emotion is then presented. Participants are asked to respond what they think the final emotion will be and to do so whenever they felt confident in their prediction. If either a response is made or 6secs have elapsed, an image of the correct emotion is presented. **(b)** Two types of trials were employed: “easy” and “ambiguous”. In easy trials, facial expressions gradually morphed towards one of the two emotions. In ambiguous trials, facial expressions remain relatively neural until two-thirds of the trial has elapsed, after which point emotion rapidly ramped up towards happy or sad.

The current study consisted of two trial types (or conditions), namely “easy” and “ambiguous”, which were modelled after previous work (Thura et al., 2012). These two trial types were interleaved throughout the runs and subjects had no knowledge of upcoming trial type. In easy trials, all intermediate faces presented were of the correct emotion (e.g., in a trial in which the correct answer is happy, no sad images are ever presented). Each successive frame had a 65% chance of being one level higher than the previous frame in the direction of the correct emotion. By the final frame, all trials had an emotion level >16. In ambiguous trials, frames within the first two-thirds of the trial (i.e., up to the 40^th^ frame) generally hovered around a neutral valence. Each successive frame had a 50% chance of being one level higher than the previous in the direction of the correct emotion and could only reach a maximum of emotion level 7. To prevent, for example, many slightly happy and a few very sad images being presented, the maximum emotion levels presented in the correct and incorrect directions were kept within two levels of each other. In the final third of the trial, there was a steep increase of emotion level in favour of the correct emotion, with a 95% chance that a given frame would be exactly one level higher than its predecessor. As with the easy condition, all trials had an emotion level >16 by the final frame.

Each subject partook in 120 trials total, divided equally across 3 blocks. Trials were evenly split between happy and sad (determined by the emotion at the final frame), and between easy and ambiguous. Trial order was randomized in every block.

### EEG Acquisition and Preprocessing

EEG was recorded continuously using a 256-channel high-impedance HydroCel Geodesic Sensory Net and the NetStation 5 acquisition software (Electrical Geodesic, Inc., Eurgene, OR). As per manufacturer standard recommendations, electrode impedance levels were kept below 50Ω during acquisition. Data were collected with a sampling rate of 1000Hz using the electrode Cz as reference with online visualization filters of 60Hz for notch, 5Hz for high-pass, and 120Hz for low-pass.

Raw data were preprocessed offline using the Automagic pipeline (Pedroni et al., 2019). First, bad channels were identified using PREP (Bigdely-Shamlo et al., 2015) in which a 1Hz high-pass filter is applied, power-line noise removed, and robust average referencing iteratively implemented to detect and interpolate bad channels to arrive at an average reference that is not affected by artifacts. Bad channels were then excluded to avoid contamination in later preprocessing steps. Second, continuous EEG recordings were filtered with a bandpass filter of 0.1-60Hz. Third, artifacts related to eye-blinks and muscle movement were corrected for using the Multiple Artifact Rejection Algorithm (MARA) – a supervised learning algorithm that uses independent component analysis to detect and eliminate artifacts based on established expert ratings (Winkler et al., 2011). Finally, the previously excluded bad channels were interpolated and the data was down-sampled to 250Hz for computational efficiency.

Data quality after preprocessing was assessed automatically by Automagic and confirmed by subsequent manual inspection. Further details regarding the Automagic pipeline can be found online (https://github.com/methlabUZH/automagic). After preprocessing and quality control, data from 57 participants were used for further analysis.

Preprocessed data files were imported into MNE-Python (Gramfort et al., 2013; Gramfort et al., 2014) for statistical analysis and visualization. Epochs were created around the stimulus onset (−1,000ms to 8,000ms time-window) and response (−1,000ms to 1,000ms time-window), with both baseline-corrected for the 500ms preceding stimulus onset. Epochs in which the activity exceeded ±150µV were excluded (average number of trials post-preprocessing: Easy=57.92±4.47, Ambiguous=57.74±4.64).

### Event Related Potentials

Here we focused on three ERPs of interest: the N170, P300 and centroparietal positivity (CPP). The N170 is a face-sensitive visually evoked ERP elicited over posterior visual cortical areas. It’s amplitude is thought to scale with how similar the stimulus is to a face (Eimer, 2011) but has also been cited as a domain-general response to unexpected perceptual events (Robinson et al., 2018). Previous work using a face-car visual discrimination task to test the drift diffusion model found that the N170 reflected an early perceptual event that is not directly related to the actual decision (Philiastides et al., 2006; Philiastides and Sajda, 2006). Based on the existing literature and on our grand-average waveform, we extracted the N170 as the peak amplitude between 120-200ms after stimulus onset at two lateral occipital sites (E114 & E168).

The centroparietal “P300” has a long-established role in in a wide variety of cognitive operations and is well defined in the literature (Sutton et al., 1965; Polich, 2012). In the current study, P300 refers to the P3b subcomponent of the P300, which has parietal topography, as opposed to the frontal P3a subcomponent. While frontal P3a represents stimulus-driven attention mechanisms, the P3b is often elicited by target detection paradigms (e.g., the oddball paradigm) though its exact role in the decision process is debated (Sutton et al., 1965; Polich, 2007). Numerous explanatory accounts have been proposed, variously implicating P3b in the allocation of attentional resources, context and memory updating, uncertainty or surprise of stimulus, and response potentiation (Hillyard et al., 1971; Polich, 2007; Nieuwenhuis et al., 2011). In one study, its magnitude covaried with the onset of a neural decision variable based on task difficulty, suggesting it may index decisional computations (Philiastides et al., 2006). In keeping with the literature and based on the maxima and time distributions observed in the grand-average waveform across our participants (Fig. 2), P300 was defined as the peak amplitude between 200-400ms after stimulus onset at electrode site Pz, as previously described (Picton, 1992).

**Fig. 2.**
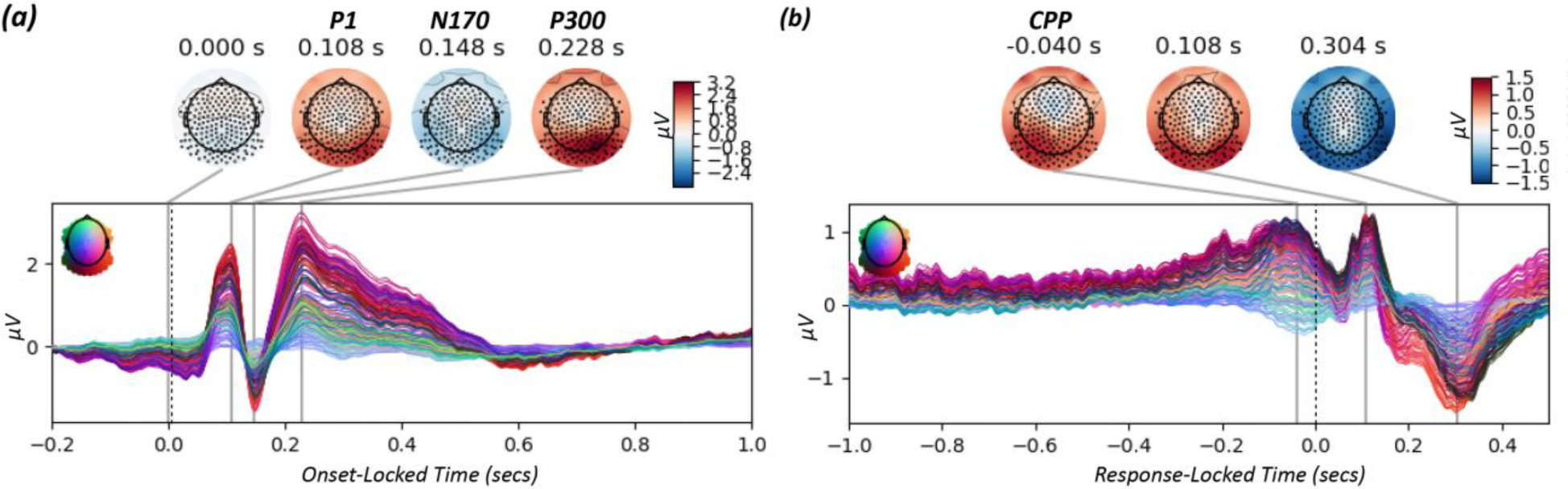
Grand average across all trials and subjects (N=57) for **(a)** onset-locked and **(b)** response-locked epochs. Line colours refer to different electrode sites as indicated in the inset. Topomaps for the timepoints of peak activity are depicted above the waveforms, with warmer and colder colours indexing higher and lower activity, respectively.

More recently, a CPP component that spatially overlaps with the P300 has also been identified and is thought to index a developing decision variable in the choice process (O’Connell et al., 2012; Kelly and O’Connell, 2013; van Vugt et al., 2019). Unlike the traditional conception of ERP components as unitary processes, the CPP is thought to be a gradual signal that scales with the strength of sensory evidence, peaking close to the time of decision/action. This rise-to-threshold like activity has been shown to be insensitive to sensory modality or target feature and unrelated to motor preparation (O’Connell et al., 2012). The CPP resembles the drift rate parameter of the DDM, which indexes sensory evidence accumulation; it is thus thought to represent decision evidence (O’Connell et al., 2012; Steinemann et al., 2018). The CPP has also been previously suggested to spatially overlap with the P3b (O’Connell et al., 2012; Twomey et al., 2015), however the two may be dissociable based on their temporal course. Unlike the P300, there is no clear established time-window of interest for CPP, with variations in the relative timing likely reflecting differences in paradigms and design (O’Connell et al., 2012; van Vugt et al., 2019). As such, we took a data-driven approach to identify a time-window of interest over which we could observe CPP signal buildup. As with O’Connell et al. (2012), we calculated the temporal slope of the activity from each participant’s average waveform at electrode site Pz in moving windows of 100ms length in 10ms steps, starting from −1,000ms to response execution (i.e., 0ms). Signal buildup rate was computed as the slope of a straight line fitted to the unfiltered signal within each sliding window. A one-tail permutation t-test implemented via *mne.stats.permutation_t_test* with 5,000 permutations was then used to identify signal buildup rates that significantly differed from 0 across all subjects in a positive direction, indicating CPP activity that is ramping up. These epochs were from 360ms before response to the time of response and are marked in black below the waveforms in Fig. 5. The cumulative sum of activity within this time-window of interest was then used to index CPP signal buildup for further analysis. An alternative to using the cumulative sum is to calculate the area under curve; however, these two signals are almost identical (Spearman’s rho=.999, *p*<.0001) and using one or the other did not affect our findings. We additionally compared our analysis to results from a larger cluster, centered on the Pz (5 electrodes: E101, E100, E129, E119, E110, E128), to ensure our findings could be replicated.

**Fig. 3.**
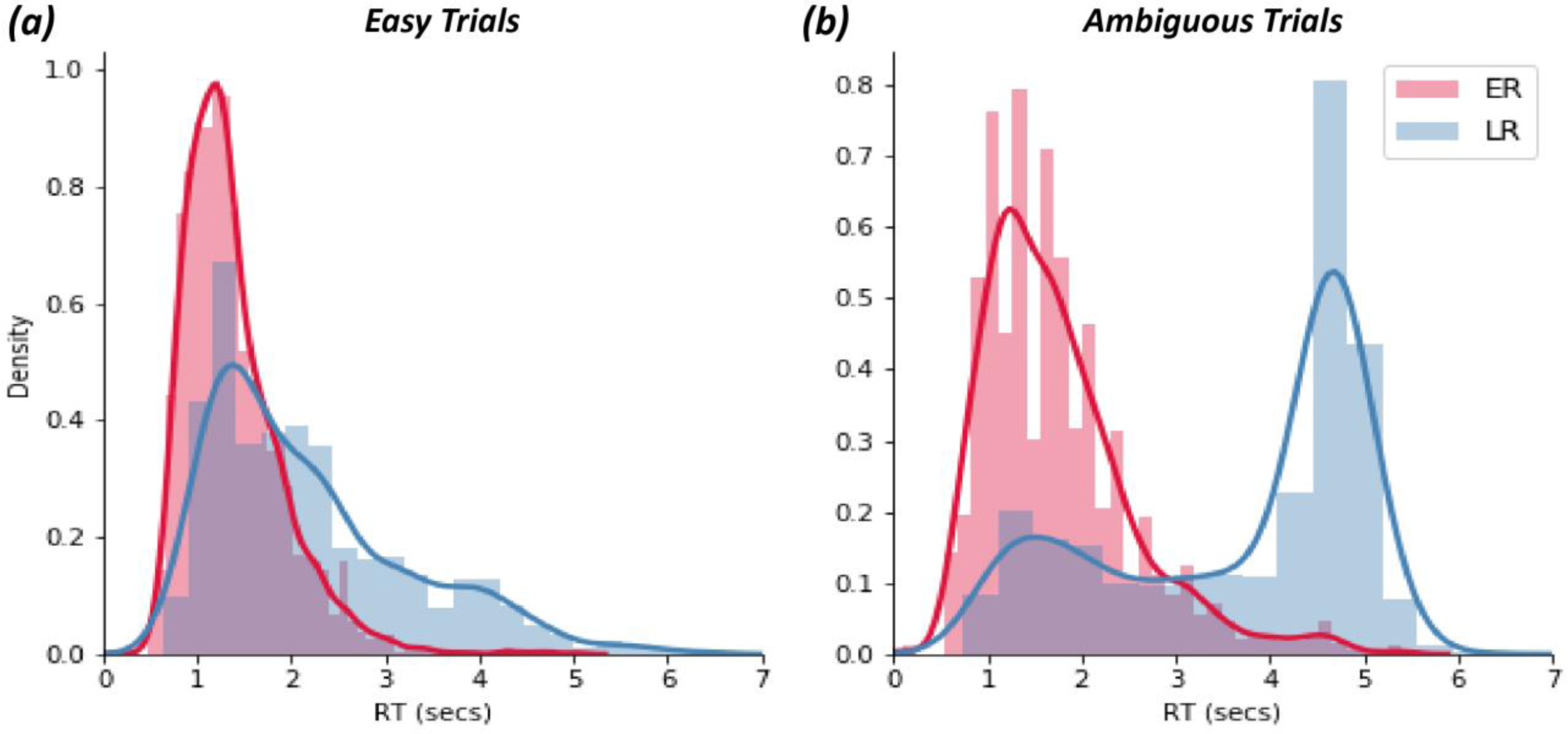
Histogram of reaction time distributions in **(a)** easy and **(b)** ambiguous trials. Solid lines reflect the gaussian kernel density estimation. ER: early responders (n=32); LR: late responders (n=25).

**Fig. 4.**
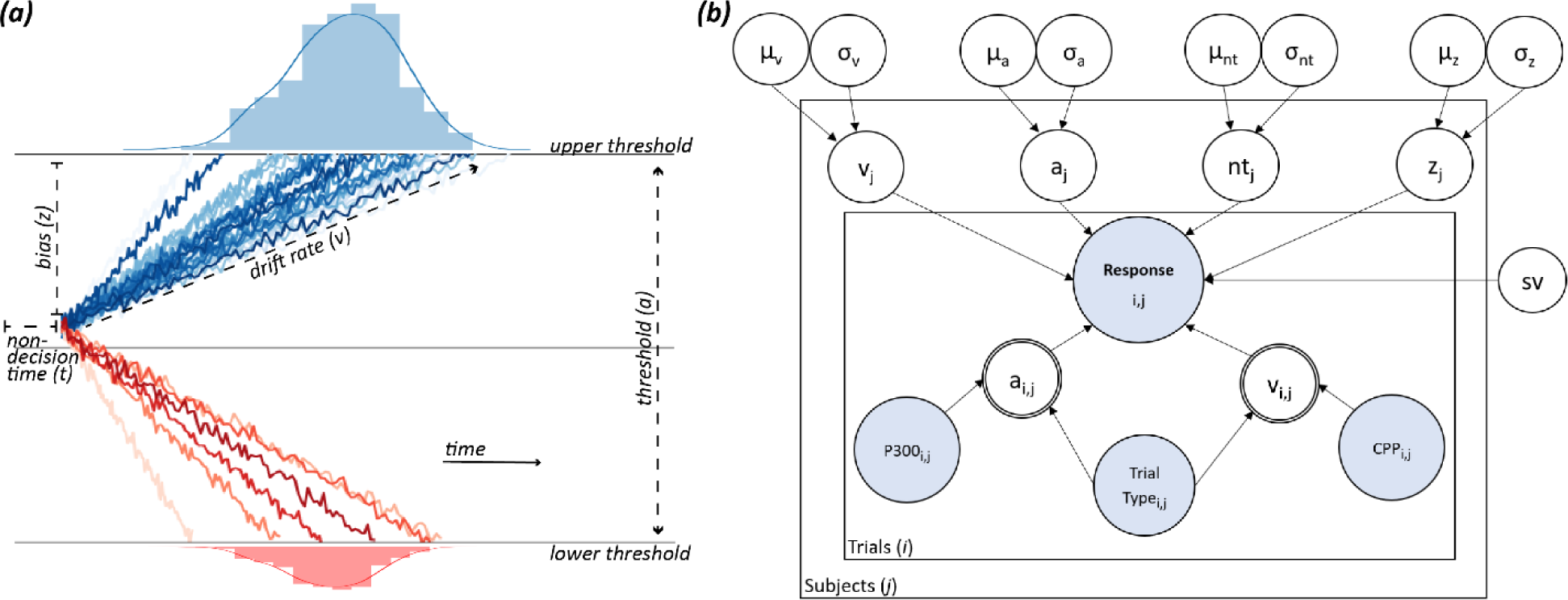
**(a)** Schematic of the drift diffusion model. **(b)** Graphical illustration of the hierarchical drift diffusion model (HDDM) with trial-by-trial neural regressors. Round nodes represent continuous random variables and double-bordered nodes represent deterministic variables, defined in terms of other variables. Decision parameters including drift rate (v), decision threshold (a), non-decision time (nt), bias(z), and standard deviation of drift rate (sv) were estimated for the group (nodes outside the plates with: group mean (μ) and variance (s)) and subjects (j) (nodes in outer plate). Blue nodes represent observed data, including trial-wise behavioral data (accuracy, RT) and neural measures (P300 and CPP). Trial-by-trial variations of v and a were modulated by P300 and CPP, respectively, as well as by trial type (i.e., easy or ambiguous trials).

**Fig. 5.**
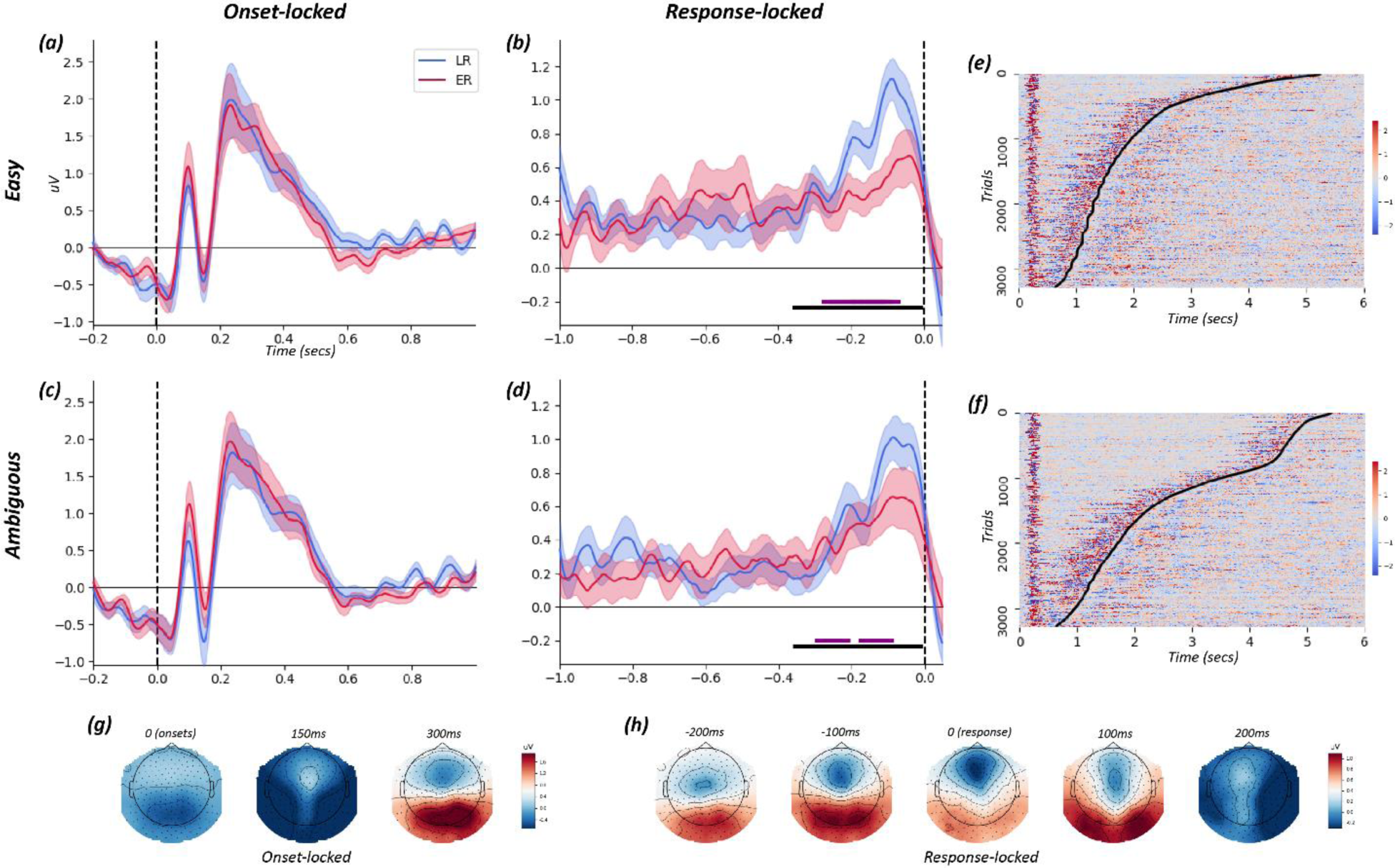
**(a-d):** Grand average waveforms from electrode site Pz. The onset-locked P300 signal is depicted in the left-hand column for **(a)** easy and **(c)** ambiguous trials. The response-locked CPP signal is depicted in the middle column for **(b)** easy and **(d)** ambiguous trials. In these CPP plots, the identified time window of interest where slopes differ from zero, indicating signal buildup, is marked by the solid black line at the bottom. Group difference in slope is marked by the solid purple line at the bottom. Blue and red lines relate to the late and early responders, respectively. Shading around the lines reflect the 95% confidence interval. **(e-f):** Single-trial plots for **(e)** easy and **(f)** ambiguous trials show the temporal relationship between the neural signal from the electrode site Pz (normalized relative to each individual’s baseline average) and decision time (curved black line). P300 can be noted early in the trial whereas CPP can be observed preceding the time of decision. **(g-h):** Topographical maps for **(g)** onset- and **(h)** response-locked activity depicted at various time points.

### Time-Frequency Analysis

In addition to standard ERP, different frequencies of oscillatory activity in the EEG signal have been linked to decision parameters of the DDM. The theta band power from mid-frontal electrodes, for example, has been linked to decision threshold and is thought to reflect a gating mechanism (Cavanagh et al., 2011a; Cavanagh et al., 2011b; Frank et al., 2015). A posterior alpha signal, argued to be harmonic to theta band activity, has also been implicated (Klimesch, 2012; Kloosterman et al., 2019) as well a motor beta signal in the contralateral hemisphere to the hand for response execution (O’Connell et al., 2012). We were interested in testing whether our straight-forward, minimalistic processing analysis of ERPs was linked to these ongoing oscillatory fluctuations. To this end, we used the Morlet wavelet methods implemented via *mne.time_frequency.tfr_morlet* to assess spectral power across the trial period on our preprocessed epoch data. The trial’s estimated power was then baseline-corrected for the 500ms preceding stimulus onset. For visualization purposes (Fig 8), the data was resampled using *scipy.signal.resample* to match onset and response across all trials (i.e., all trials began at timepoint 0 and ended at timepoint 1 which stood for the maximum RT for the subject). Power was averaged across the frequency range for each band (alpha=0-4Hz, theta=4-8Hz, alpha=8-12Hz, beta=12-30Hz, gamma=30-45Hz). Each sample of EEG time course was z-scored and outliers (*z*>4.5) were replaced with the average EEG power (Frank et al., 2015). We then extracted the early (first 10% of trial time after onset) and late (last 10% of trial time before response) power for each frequency band to compare against ERPs of interest. Based on the cited literature, we focused on theta power from the FCz, alpha from the Pz, and beta from electrodes of the left hemisphere. We also repeated this analysis using the power for each frequency band averaged across all electrodes.

**Fig. 6.**
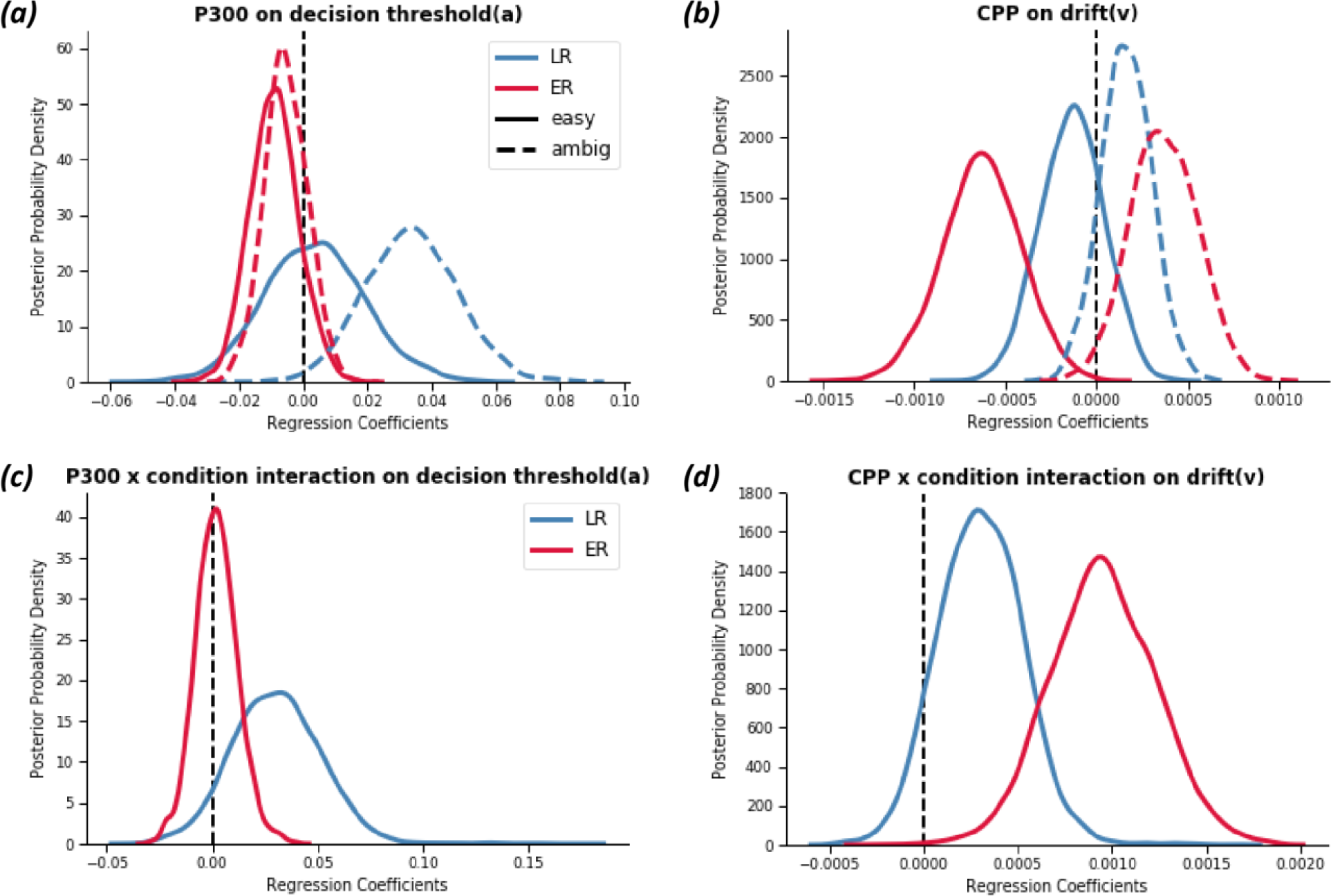
Bayesian posterior probability densities for modulation of decision parameters estimated from the hierarchical drift diffusion model by neural signals. Peaks reflect the best estimates, while width represent uncertainty. Simple effects of **(a)** P300 on decision threshold and **(b)** CPP on drift rate are depicted in the upper row. Interaction effect of **(c)** P300 **(d)** and CPP on decision threshold and drift rate, respectively, with condition are depicted in the lower row. A more positive regression coefficient indicates ambiguous > easy.

**Fig. 7.**
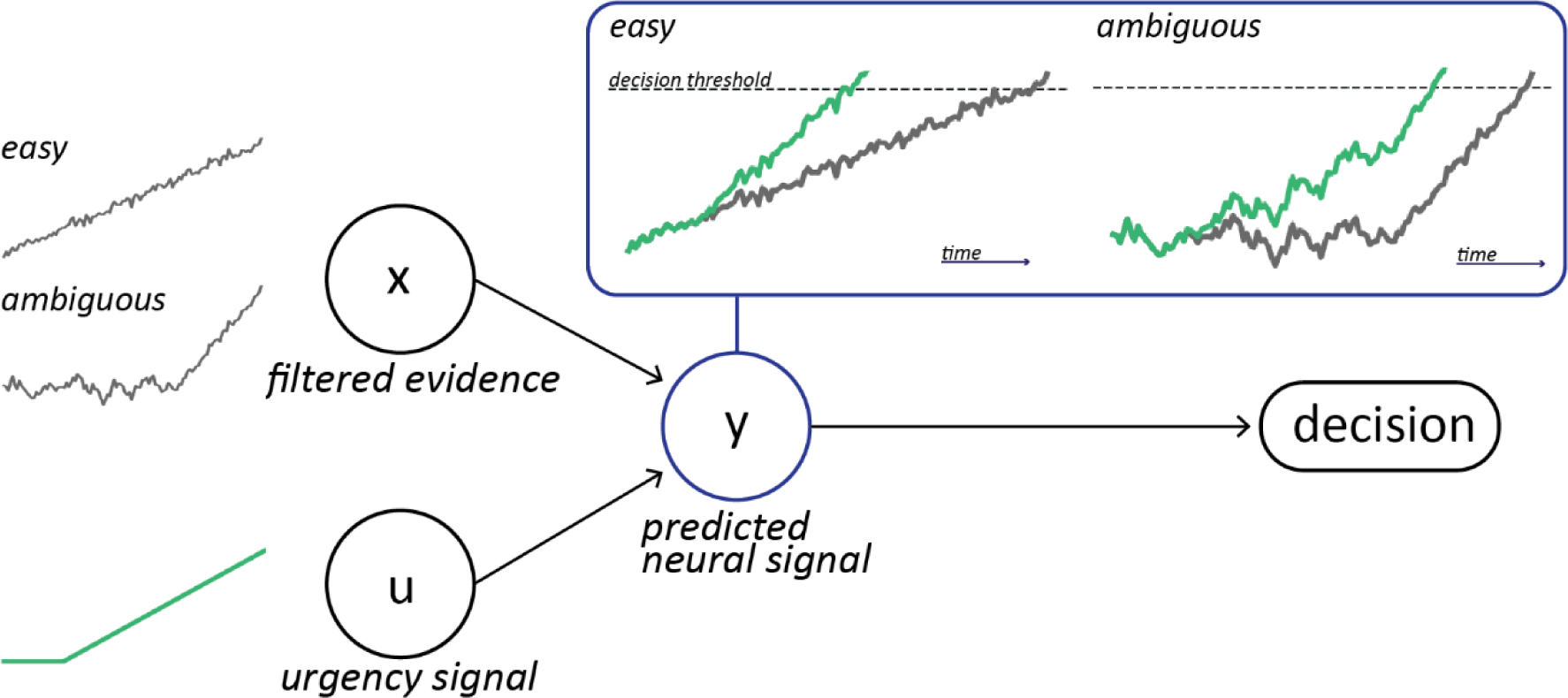
Schematic of the urgency gating model. Sensory evidence is first differentiated and filtered. The resulting signal (x) is then multiplied by a subject’s evidence-independent urgency signal (u) that grows in time. The combined signal together forms the model’s predicted neural signal (y). Green lines depict a neural signal that incorporates an urgency signal whereas grey lines do not. Once the predicted neural signal crosses a decision threshold, a decision is made.

**Fig. 8.**
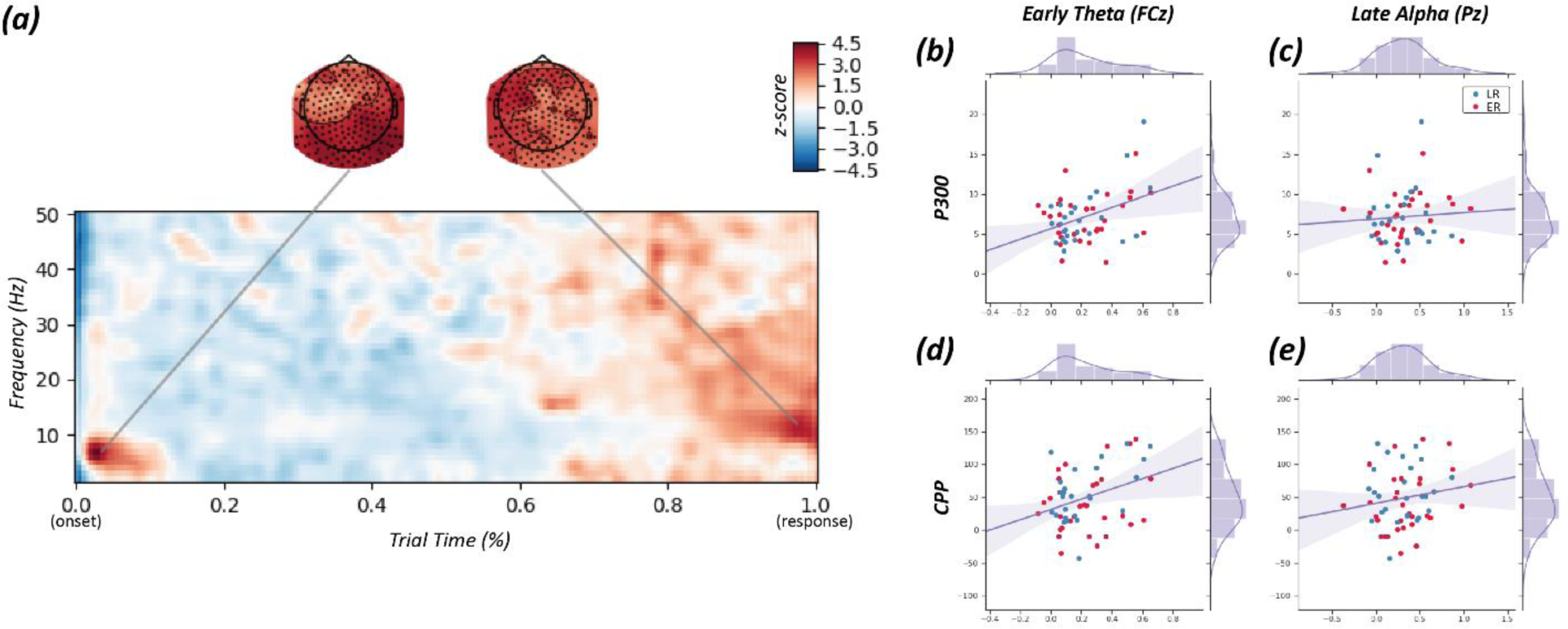
**(a)** Average time-frequency power across all trials. Data is resampled to match onset (timepoint 0) and response (timepoint 1). A strong early theta power and a late alpha power can be observed. Heatmap depicts strength of power, as compared to baseline (−500ms to onset), with warmer and colder colours reflecting higher or lower power, respectively. **(b-e)** Scatterplots with regression lines depicting correlation between ERPs of interest (i.e., P300 and CPP) and EEG oscillatory power (i.e., early theta and late alpha).

### Tendency to Wait: Early versus Late Responders

As with our previous work (Yau et al., 2020), we observed two distinct groups of individuals based on performance under the ambiguous condition (Fig. 3) : (1) early responders (*n*=32), who on >=80% of ambiguous trials responded during the first two-thirds of the trial before information ramped towards one direction and (2) the rest, who were categorized as late responders (*n*=25). We hypothesized that one factor that may drive this group difference lies in differences in an endogenous urgency signal.

One mixed-design ANOVA per neural signal (i.e., N170 maximum amplitude, P300 maximum amplitude, and CPP cumulative sum) was conducted to investigate the within-subject relationships of conditions (i.e., easy and ambiguous) and between-subject relationships for group affiliation (i.e., early and late responders) as well as their interaction. Given the unequal sample size, Levene’s test was used to test and confirm equality of variance between the two groups. If sphericity was violated, Greenhouse-Geisser corrected degrees of freedom are reported.

In addition, given that CPP gradually ramps up in time, we tested whether CPP buildup (slope) may differ between the two groups at specific time intervals within our larger time-window of interest. A two-tailed, two sample permutation t-test (*mlxtend.evaluate.permutation_test*) with 5,000 permutations was conducted per condition. Time windows where the two groups significantly differed are marked in purple below the waveforms in Fig. 5.

### Hierarchical Drift Diffusion Model

Drift diffusion models (DDM) are commonly used to infer latent processes underlying perceptual decision-making and to link them to neural mechanisms (Ratcliff et al., 2016). In the DDM framework, decision-making between two alternatives is reflected by a continuous integration of sensory evidence over time until a decision threshold for one of the choices is reached. The model decomposes behavioral data into four parameters: non-decision time (*nt*) for stimulus encoding and response execution latencies, bias (*z*) towards one choice alternative, drift rate (*v*) for speed of evidence accumulation, and decision threshold (*a*) which determines how much evidence is needed before a decision is made (Fig. 4a). The shape of the reaction time (RT) distribution determines the decision parameters (Ratcliff et al., 2016).

Here, we applied a hierarchical estimation of the DDM (HDDM) (Wiecki et al., 2013), implemented in Python 2.7 (http://www.python.org), to calculate the decision parameters (Fig. 4b). The hierarchical design assumes that model parameters from individual participants, while varying, are not completely independent. Rather, individuals’ parameters are drawn and constrained by priors based on the group distributions (Gelman et al., 2013). This Bayesian estimation is thought to be more robust in recovering model parameters, particularly when the number of trials is relatively small (Matzke and Wagenmakers, 2009; Wiecki et al., 2013). Trials that fell within 5% of each tail of the RT distribution were considered outliers that cannot be captured by HDDM (e.g., slow responses due to inaction or fast erroneous responses due to action slips) and removed from analysis (Wiecki et al., 2013). Markov chain Monte Carlo sampling was used for Bayesian approximation of the posterior distribution of model parameters. 5,000 samples were drawn from the posterior to obtain smooth parameter estimates, while the first 100 samples were discarded as burn-in. Convergence of Markov chains were assessed by inspecting traces of model parameters, their autocorrelation, and computing the Gelman-Rubin statistic (Gelman and Rubin, 1992) to ensure that the models had properly converged.

As a first step, we constructed a base model whereby decision parameters are simply a function of RT and accuracy. In a second model, we expanded upon this base with a simple model that allowed drift rate and decision threshold to vary between easy and ambiguous conditions. Our third model further extended this by allowing for trial-by-trial variations in neural activity, in addition to condition, to modulate decision parameters. The estimated posterior distributions index the degree to which the decision threshold (*a*) is altered by variations in P300 and N170 and how the drift rate (*v*) is explained by CPP buildup. Our comprehensive model is as follows: [*a*(*t*) *=* β_0_ + β_1_P300(*t*) * β_2_N170(*t*)*β_3_condition(*t*), *v*(*t*) *=* β_4_ + β_5_ CPP(*t*)*β_6_condition(*t*)]. In these regressions, a larger positive coefficient weight (β) indicates a stronger positive correlation between neural measure and decision parameter, and *vice versa*. Of note, the decision threshold and drift rate parameters were estimated separately for early and late responders for all models using the “*depends_on”* function in HDDM, as the two groups are assumed to have different parameter distributions. Further, we iteratively added in modulators to test whether they improved model fit (described below).

The deviance information criterion (DIC) was used for model comparison (Spiegelhalter et al., 2002). A lower raw DIC value for a given model (for the whole group) favors models with highest likelihood and least number of parameters. A DIC difference of 10 is considered significant (Zhang and Rowe, 2014). All reported DIC values are relative to the base model (i.e., target model DIC minus base model DIC) – the more negative the value, the better the model fit compared to the base model. Parameters of the best fitting model were analyzed by Bayesian hypothesis testing which examines the probability mass of the parameter region in question (i.e., percentage of posterior samples greater/smaller than zero). For all HDDM analyses, we considered a posterior probability ≥95% of the respective parameters being different than zero as significant (Wiecki et al., 2013).

### Urgency Gating Model

Models of decision-making incorporating an endogenous urgency signal posit that choices result from a combination of signals that reflect the available sensory evidence and an evidence-independent urgency signal that grows in time (Cisek et al., 2009; Drugowitsch et al., 2012). We constructed a minimalistic urgency gating model (UGM) and an urgency-free, non-hierarchical drift diffusion model (DDM) to compare against and to test whether accounting for an urgency signal may better fit our observed data.

In both models, a filtered evidence variable *x* was derived by the following differential equation:

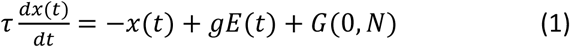

At any given time *t*, the evidence *E* (i.e., level of information/facial emotion level) is multiplied by an attentional fixed gain term *g*. An intra-trial Gaussian noise variable *G*(0,*N*) with a mean of 0 and a standard deviation of *N* was added. A *N* of 6 was chosen based on previous work (Carland et al., 2015; Yau et al., 2020) and because it gave a range of simulated RTs with similar variability as the observed data in our current study. The time constant *τ* determines how far back in time sensory information is considered by the model. The UGM posits that only recent information is used to inform decision whereas the DDM does not; thus, *τ* was set to 200ms for the UGM on the basis of previous behavioral and physiological studies (Cisek et al., 2009; Thura et al., 2012; Thura and Cisek, 2014) while the maximum trial duration of 6sec was used as *τ* for the DDM. Evidence (*E*), gain (*g*), and noise (*N*) parameters were the same in both models.

Next, the filtered evidence *x* at a given time t was used to compute the estimated neural activity *y* as follows:

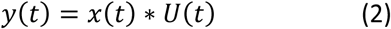

Where *U*(*t*) = *u* · *t* and represents the urgency signal that rises from zero with a slope *u*. The UGM assumes that evidence is multiplied by the urgency signal which increases linearly with time (Cisek et al., 2009; Thura and Cisek, 2014; Yau et al., 2020). An alternative non-linear model fitted with an additional 2^nd^ order polynomial parameter did not significantly improve model fit, as such, we opted to keep the linear model with the minimum number of parameters. A decision is made when the variable *y*(*t*) reaches a critical decision threshold *α*. A core prediction of the UGM is that decisions made with low levels of filtered evidence *x* should be associated with high levels of urgency and *vice versa*. In other words, high urgency will push the individual to commit to a choice even if evidence for that choice is weak. On the other hand, the DDM does not have an urgency signal and *U*(*t*) = 1; thus, a decision is made only when the variable *x*(*t*) reaches threshold *T*. In both models, a non-decision time of 200ms was added (Yau et al., 2020).

Each model adjusts for one parameter: for UGM, the *u* parameter and for DDM, the *α* parameter. Both of these parameters influence the means of RT distributions. An exhaustive search was implemented to find the parameter value that minimized the mean squared error between each model’s predicted RT and the observed RT across all trials for a subject. The models were used to simulate 5,000 trials, the mean RT was used to compare against the real RT distributions.

Linear mixed effect models implemented via *statsmodels.mixedlm* were used to examine the relationship between the observed CPP signal compared to a neural code predicted by either the UGM or the DDM at the trial-level. Subjects were included as a random variable with differing intercepts. For both the predicted and observed neural signal, buildup was determined as the cumulative sum of activity from 500ms post stimulus-onset to the time of response. This 500ms delay was to ensure that we did not confound CPP activity with the P300 signal, as they overlap spatially. The log-transformed absolute value of the predicted neural code was used since CPP is assumed to be a general evidence accumulation signal that positively ramps up regardless of the stimulus presented. As with the HDDM, trials in which participants registered no response or with RT that fell within 5% of each tail of the RT distribution were considered outliers and discarded.

### Statistical Analysis of Behavioral Data

Statistical tests were conducted using packages *pingouin (Vallat, 2018), statsmodels* (Seabold and Perktold, 2010), and *mlxtend* (Raschka, 2018) in Python 3.7 (http://www.python.org). To account for potential spurious outliers in our relatively low sample size, non-parametric tests were used to assess subject-level data. Mann-Whitney U tests were conducted to compare differences in behavioral performance (i.e., mean accuracy and RT) between emotion, groups, as well as neural signals between correct and incorrect trials. Spearman correlations were used to examine relationships between different neural signals (e.g., P300 and CPP) and other metrics of interest (e.g., HDDM decision threshold, DDM decision threshold, and UGM urgency signal). An alpha of .05 was used as the threshold for statistical significance and results were corrected for multiple comparisons using the FDR Benjamini-Hochberg correction as implemented by *statsmodels.stats.multitest.multipletests*.

## Results

### Behavioral Results

The early responder (ER) group were significantly less accurate on ambiguous trials (mean=53.76%±6.23) than the late responder (LR) group (mean=73.36±7.34) (*U*=629.5, *p<*.0001). ER individuals appear to be performing at chance level, suggesting that they were guessing rather than making informed judgements. Difference in accuracy performance was also observed on easy trials with the ER group (mean=94.9%±5.42) having lower accuracy than the LR group (mean=98.9%±1.2) (*U*=766.0, *p*<.0001) although both groups preformed near ceiling. Moreover, although group categorization was made based on RTs on ambiguous trials (see *Methods*), ER (mean=1.39s±0.28) tended to also respond earlier than LR (mean=2.16 ±0.61) on easy trials (*U*=718.0, *p*<.0001). Overall, subjects were neither faster (*U*=6165.5, *p*=.498) nor more accurate (*U*=5536.0, *p*=.107) to either happy or sad stimuli.

### Comparing Neural Signals of Interest between Early and Late Responders

To examine endogenous determinants of decision RT, we began by identifying the neural correlates previously implicated in evidence accumulation during perceptual decision-making. In addition to a prominent sensory-evoked P300 (Fig. 2), we observed a CPP activity that increased over time and peaked close to the time of response – consistent with the build-to-threshold dynamics proposed by drift diffusion models. These two signals were significantly positively correlated (*rho*=.38, *p*<.0001). The grand-average waveforms indicate that CPP generally peaked before response suggesting that CPP encodes sensory information and not motor readiness, as shown by others (O’Connell et al., 2012; Kelly and O’Connell, 2013). Crucially, the use of a gradual morphing stimuli eliminated sensory-evoked deflections (e.g., N170 and P300) from the ERP trace time-locked to the response, making it possible to disentangle and finely trace the evolution of the CPP from its onset to its peak. Given the temporal evolution of the CPP, we probed with a permutation test whether there may be windows of time within which the buildup rate differs between ER and LR groups. For easy trials, we found that LRs had greater CPP signal buildup compared to ERs during the −280ms to −64ms preceding response (Fig. 5b). For ambiguous trials, we again observed greater CPP signal buildup among LRs than ERs over two time-windows preceding response: −300ms to −204ms and −180ms to −84ms (Fig. 5d). This supports the premise that CPP reflects the accumulating decision evidence.

We next sought to examine whether N170 amplitude, P300 amplitude, and CPP within a broader time window of interest (*see Methods*) may differ between groups (ER vs LR) or between conditions (easy vs ambiguous) (Fig. 5). Contrary to our hypothesis, no significant main effect of group (*F*(1,55)=2.409, *p*=.126), condition (*F*(1,55)=.059, *p*=.809), or their interaction (*F*(1,55)=0.113, *p*=.738) was observed for the N170. For P300, no significant main effect of group (*F*(1,55)=0, *p*=.986), condition (*F*(1,55)=0, *p*=1.00), or their interaction (*F*(1,55)=0.718, *p*=.402) was observed. Similarly for the CPP, no main effect of group (*F*(1,55)=1.708, *p*=.197), condition (*F*(1,55)=1.611, *p*=.210), or their interaction (*F*(1,55)=2.399, *p*=.127) was statistically significant. Additionally, N170 amplitude (*U*=1631, *p*=.973), N170 latency (*U*=1794, *p*=.845), P300 amplitude (*U*=1646, *p*=.973), P300 latency (*U*=1649, *p*=.973), and CPP (*U*=1862, *p*=.845) did not differ between correct and incorrect trials – suggesting these signals may not only reflect external evidence but also an internal decision quantity as responses were nonetheless made despite incorrect trials being more likely to occur when external sensory evidence is ambiguous. In other words, these ERPs are more closely associated with the choice that was made (i.e., internal estimate of evidence) rather than the external sensory evidence.

### Trial-by-trial Variations in Neural Signals Modulates Decision Parameters

Though we did not observe discernable differences when examining neural signals alone, one might nonetheless postulate that these signals may relate differently to decision parameters depending on group affiliation and condition. We thus assessed whether decision parameters, as estimated by the HDDM, are modulated by EEG neural signals. To this end, we examined the relationship between the decision parameters and neural signals at a trial-by-trial level to estimate their regression coefficient. Non-neural HDDM models where both decision threshold and drift rate varied as a function of condition had improved model fit (difference in DIC=-1262.89) compared with just threshold (DIC=-86.1) or just drift rate (DIC=-1236.46). Moreover, allowing the decision threshold and drift rate to vary parametrically with P300 amplitude and CPP buildup, respectively, yielded a better fitting model (DIC=-1272.87) compared to just P300 (DIC=-86.15), CPP (DIC=-1243.81), or no neural signal (DIC=-1236.46). Allowing decision threshold to vary with N170 did not improve model fit (DIC=87.97), suggesting it does not reflect the decision evidence used – at least in the context of the DDM. In sum, our best fitting model was one that allowed for decision threshold and drift to vary by condition, with the P300 influencing threshold and the CPP drift rate, respectively.

Trial-by-trial modulation of P300 amplitude (Fig 6a & c) was parametrically related to higher decision thresholds, but only in LRs and only for ambiguous trials (99.20% posterior probability > 0). This is confirmed in the interaction analysis; P300 was significantly higher on ambiguous compared to easy trials among LRs (95.33% posterior probability > 0). Regression coefficients indicate all other relationships were not significant. This is in keeping with the P300 reflecting sensory evidence accumulation, as a higher decision threshold in longer (ambiguous) trials would lead to increased area under the curve of the evidence and hence greater EEG signal (Kelly and O’Connell, 2015). Our results suggest that, in LRs, the P300 may therefore reflect the effect of a “caution” or “inverse-urgency” signal, i.e. elevated decision threshold, when the environment lacks strong information for one choice over the other. In ERs, however, the relationship between P300 and decision threshold is not seen, probably due to the short latency to respond.

CPP, as explained above, is thought to index the time-varying amount of information available, and thereby the drift rate (Fig 6b & d). Among ERs, CPP was related to lower drift rate on easy trials (99.99% posterior probability < 0), likely because these individuals still tend to respond relatively early on easy trials and at points of low level of information. However, on ambiguous trials, CPP was in fact related to higher drift rate in ERs (97.98% posterior probability > 0). Interaction analyses suggest that CPP was significantly related to greater increases in drift rate in ambiguous compared to easy trials among ERs (99.90% posterior probability > 0) and a similar trend was noted among LRs (91.41% posterior probability > 0). This supports the notion that CPP indexes an accumulating decision variable. Moreover, given that ERs perform around chance on ambiguous trials but still demonstrate high levels of CPP signal buildup suggests that CPP may also index an alternative parameter no captured by the DDM which drives decision-making, even when the actual evidence is ambiguous.

### CPP Reflects a Combination of Evidence Accumulation and an Evidence-Independent Urgency Signal

We hypothesized that the CPP signal buildup across the entirety of a trial reflects a combination of both the sensory evidence available in the environment and an evidence-independent urgency signal. As discernable from the waveform plot (Fig. 5d), CPP nonetheless ramps in time among ERs despite their decisions often being made in situations of ambiguous sensory evidence. This raises the possibility that CPP reflects a multiplicative effect of evidence and urgency as in eq. (2). We therefore tested a second model that incorporates an urgency signal (Fig. 7; described in *Methods*). As expected, the estimated urgency signal was significantly lower for the LR (mean=1.73±1.27) compared to the ER (mean=6.60±3.39) group (*U*=26.5, *p*<.0001).

The predicted neural signal from a model that included the evidence-independent urgency signal significantly related to our actual recorded CPP signal (β=43.74, *p*=.023, 95%CI=[6.17, 81.33]). We did not observe this with the predicted neural signal from the DDM (β=7.155, *p*=.706, 95%CI=[-30.07, 44.38]), without the urgency signal. A pairwise Spearman correlation indicates that our DDM and HDDM yielded highly similar decision threshold estimates (*rho*=.951, *p*<.0001) and are, therefore, comparable. Taken together, our findings confirm that the observed CPP signal fits with predictions of the UGM (eq. 2 and Fig. 7).

### Relationship between ERPs and Oscillatory Fluctuations in EEG Signals

Difference frequencies of EEG oscillations have been previously implicated in perceptual decisions (e.g., Cavanagh et al., 2011a; Klimesch, 2012). Here, we aimed to link our ERPs to these bands of EEG oscillations. Our paradigm allowed us to identify when in the decision-process this signal peaked. We observed an early theta power near stimulus onset and a late alpha power close to the time of response. Pairwise Spearman correlations indicate that early mid-frontal theta power was related positively to both P300 maximum amplitude (*rho=*.329, *p*=.012) and CPP buildup (*rho*=.411, *p*=.001). However, late posterior alpha did not relate significantly to either P300 (*rho*=.116, *p*.389) or CPP (*rho*=.147, *p*=.274). No other significant correlation was observed between either the early or late power of other frequency bands and our ERPs of interest.

## Discussion

The current study interrogates the neural determinants of perceptual decision-making in humans by isolating discrete EEG signatures associated with the parameters of a computational model. We used ERP to identify correlates of the sensory evidence and decision variable (Gold and Shadlen, 2007; Kelly and O’Connell, 2015), and urgency signaling (Cisek et al., 2009; Thura et al., 2012). By using a slowly morphing facial stimulus with variable transition rates, we were able to dissociate a sensory evidence accumulation component (the P300, time-locked to stimulus onset) and a decision variable (the CPP, time-locked to the choice). This allowed us to link the CPP to an accumulating decision variable by showing that it builds up to the time point of the decision and is proportional to the drift rate from the DDM. Moreover, we demonstrate that CPP best reflects the product of evidence and a linear urgency signal as predicted by the urgency gating model and is not accounted for by the DDM.

We tested the urgency gating model by using a slowly morphing stimulus with ambiguous trials, in which there was no evidence early on favoring one response over the other (Cisek et al., 2009). As in our previous work (Yau et al., 2020), our volunteer cohort spontaneously separated into early and late responders. The early responder group consistently responded faster, even in easy trials, and performed at chance in ambiguous trials as they responded before the facial emotion information had appeared. Nonetheless, the early responders had evidence of ramping CPP activity, which can best be explained by the presence of a multiplicative urgency signal driving the early decision. This endogenous urgency signal varied between individuals, suggesting that it may be thought of as a cognitive trait (Carland et al., 2019).

We found that the domain-general CPP signal gradually builds throughout the trial, ramping up steeply close to decision, and resembled characteristics of a neural signature of decision formation. As with previous studies (Kelly and O’Connell, 2013; van Vugt et al., 2019), the peak of CPP activity temporally preceded that of the response, suggesting it reflects an intermediate level in the decision hierarchy between stimulus onset and motor action. Further supporting the role of CPP in evidence accumulation, we found that its amplitude covaried with the drift rate parameter from the DDM, though this relationship was context-dependent and modulated by individual differences in tendencies to wait. Importantly, our results challenge the notion that CPP solely traces sensory evidence (O’Connell et al., 2012) in two key ways: (i) CPP was related to higher drift rate in situations of high, compared to low, ambiguity in the decision environment, and (ii) the group of subjects who tended to respond early in the trial when sensory evidence is low, and performed around chance, still demonstrated CPP signal buildup. These results identify the CPP as a decision variable, and dovetail with recent findings that CPP is mediated by subjective evidence and perceived decision confidence, over and above the sensory evidence (Herding et al., 2019; Tagliabue et al., 2019). Although we cannot identify the source of the CPP, a combined fMRI-EEG study found the same posterior distribution of the CPP shown here (Fig. 5h) and localized its origin to the supplementary motor area (Pisauro et al., 2017).

Our findings lend support to the consideration that the decision variable reflects a combination of sensory evidence and a dynamic urgency signal that pushes one to commit to a choice even if sensory evidence is weak (Cisek et al., 2009). It must be noted that the evidence-independent urgency signals could be misconstrued as drift rate or threshold effects in pure evidence accumulation models; to disambiguate the two, one needs to dynamically manipulate the amount of information presented (Thura et al., 2012), as in the present study. Indeed, neural signals predicted by a model that accounts for an individual’s urgency signal fit better with our observed CPP signal than that predicted by the conventional drift diffusion model. Note that this urgency signal was estimated per subject and reflects a global mechanism affecting decision-making that is not thought to be specific to any one sensory modality. Such a global gain modulation may not only manifest in the firing rate of neurons tracking the evolving decision process; urgency may influence processes both early and late in the decision hierarchy such as in the gain of sensory inputs to decision circuits (Heitz and Schall, 2013) and in downstream regions involved directly with motor execution (Thura and Cisek, 2016, 2017; Steinemann et al., 2018).

Finally, the use of relatively long trial times with smooth sensory transitions allowed us to temporally disentangle the CPP from it’s spatially overlapping counterpart, the P300. The P300 has been frequently implicated in decision-making since it’s discovery (Sutton et al., 1965) and several lines of evidence have converged to show that the P300 component serves as a bridging step between stimulus processing and response preparation (Donchin and Coles, 1988; Polich, 2007; San Martín et al., 2013), though there is little consensus regarding its precise functional role. Here, we demonstrated by onset-locking to the visual stimulus that P300 was related to increased decision thresholds only in individuals tending to wait when information is ambiguous. This suggests that the P300 is under the influence of a caution signal setting a higher decision threshold in late responders. One prominent theory on the biological origins of P300 amplitude is that it is a cortical manifestation of the phasic locus-coeruleus-noradrenergic orientation response which potentiates information processing and prepares or facilitates a behavioral response to the eliciting stimulus (Swick et al., 1994; Nieuwenhuis et al., 2011). This may underpin the famous sensitivity of the P300 to stimulus probability (Mars et al., 2008; Lucci et al., 2016) and motor inhibition (Smith et al., 2008) – concepts which, in the terminology of the evidence accumulation framework, translate to changes in the decision threshold. Unlike the P300, the N170, which is also observable by time-locking the EEG signal to the onset of face presentation, is linked to early encoding of facial features and not thought to encode higher level features such as emotion (Eimer, 2011); as such, it did not correlate with any parameters of the DDM. Collectively, our findings suggest that the P300 and CPP are dissociable and play different roles in the decision process. The short trial time implemented in previous studies may have led to the theory that they are one and the same (Verleger et al., 2005; O’Connell et al., 2012; Twomey et al., 2015).

Although the notion of urgency in decision-making has been gaining momentum, there remains debate how a hypothetical urgency signal is incorporated into the decision process and where it originates. One potential alternative interpretation of our findings is that decisions results from boundary adjustments over the course of a trial. Previous psychophysical studies in humans have found collapsing bound to improve model fit (Tajima et al., 2016; Palestro et al., 2018), though negative findings exists (Hawkins et al., 2015; Voskuilen et al., 2016). This may be a matter of interpretation: an increasing urgency signal is mathematically equivalent to a symmetrically collapsing decision threshold. However, as with other studies investigating the neural signals of urgency (Cisek et al., 2009; Thura et al., 2012), results from our study indicates that urgency works via a dynamic gain in evidence accumulation that increases with elapsing time. For simplicity, we utilized a minimalistic UGM (equation 2) to make it comparable with the DDM; however, it is possible that the baseline or starting point of the dynamic gain is non-zero and can be an additional parameter to fit (Thura et al., 2014; Trueblood et al., 2020). Nonetheless, open questions remain regarding where urgency might be generated in the brain. According to the affordance competition model, the basal ganglia are thought to bias decisions via cortico-striatal connections (Cisek, 2007) and neural recordings in monkeys suggest that the urgency signal comes through the output nuclei of the basal ganglia (Thura and Cisek, 2017). In agreement with this proposal, preliminary findings in humans point to the caudate as a potential root of the urgency signal in our task (Yau et al., 2020). However, there are other brain regions that could also encode urgency, for example, the locus coeruleus (Murphy et al., 2016). Further study is warranted to better identify the source of the urgency signal in the brain.

## Conclusion

Our results reveal how different decision parameters may be reflected in neural signals. In particular, we demonstrate that the CPP, which behaves as a developing decision variable, is a reflection of the sensory evidence available in the decision environment combined with an endogenous urgency signal that grows in time. By embedding these neural signals into a computational framework, it is possible to generate testable predictions about how different parameters should vary as a function of specific stimulus properties such as discriminability. These mechanisms expose principles of cognitive function in general and can pave a new and more precise understanding of how clinical brain disorders and experimental manipulations impact on decision-making in the human brain.

## Acknowledgements

This research was supported by grants from the Canadian Institutes of Health Research and the Natural Sciences and Engineering Research Council of Canada to AD. YY is a Vanier Scholar and received funding from the Canadian Institute of Health Research. YY and AD designed research; YY and MT collected the data; YY and TH analyzed the data and contributed methods; and YY, TH, PC, LF, and AD contributed to result interpretation. YY drafted the initial manuscript; all authors contributed to writing of this manuscript. We would like to thank Michael J Frank and his group on advice for using the hierarchical drift diffusion model.

